# Lancet: genome-wide somatic variant calling using localized colored DeBruijn graphs

**DOI:** 10.1101/196311

**Authors:** Giuseppe Narzisi, André Corvelo, Kanika Arora, Ewa A. Bergmann, Minita Shah, Rajeeva Musunuri, Anne-Katrin Emde, Nicolas Robine, Vladimir Vacic, Michael C. Zody

## Abstract

Reliable detection of somatic variations is of critical importance in cancer research. Lancet is an accurate and sensitive somatic variant caller which detects SNVs and indels by jointly analyzing reads from tumor and matched normal samples using colored DeBruijn graphs. Extensive experimental comparison on synthetic and real whole-genome sequencing datasets demonstrates that Lancet has better accuracy, especially for indel detection, than widely used somatic callers, such as MuTect, MuTect2, LoFreq, Strelka, and Strelka2. Lancet features a reliable variant scoring system which is essential for variant prioritization and detects low frequency mutations without sacrificing the sensitivity to call longer insertions and deletions empowered by the local assembly engine. In addition to genome-wide analysis, Lancet allows inspection of somatic variants in graph space, which augments the traditional read alignment visualization to help confirm a variant of interest. Lancet is available as an open-source program at https://github.com/nygenome/lancet.

---

Reliable detection of somatic variants from next-generation sequencing data requires the ability to effectively handle a broad range of diverse conditions such as aneuploidy, clonality, and purity of the input tumor material. The sensitivity and specificity of any somatic mutation calling approach varies along the genome due to differences in sequencing read depths, error rates, mutation types and their sizes (e.g., SNVs, indels, CNVs). Micro-assembly approaches^1^ have been successful at calling indels up to a few hundred base pairs in length, allowing inquiry into the twilight zone between longer indels and shorter CNVs. However, existing micro-assembly methods rely on separate assembly of tumor and matched normal data, which has limitations in regions with low supporting coverage, repeats, and large indels. Accounting for these variables requires flexible methods that can adapt to the specific context of each genomic region.

**Figure 1.**
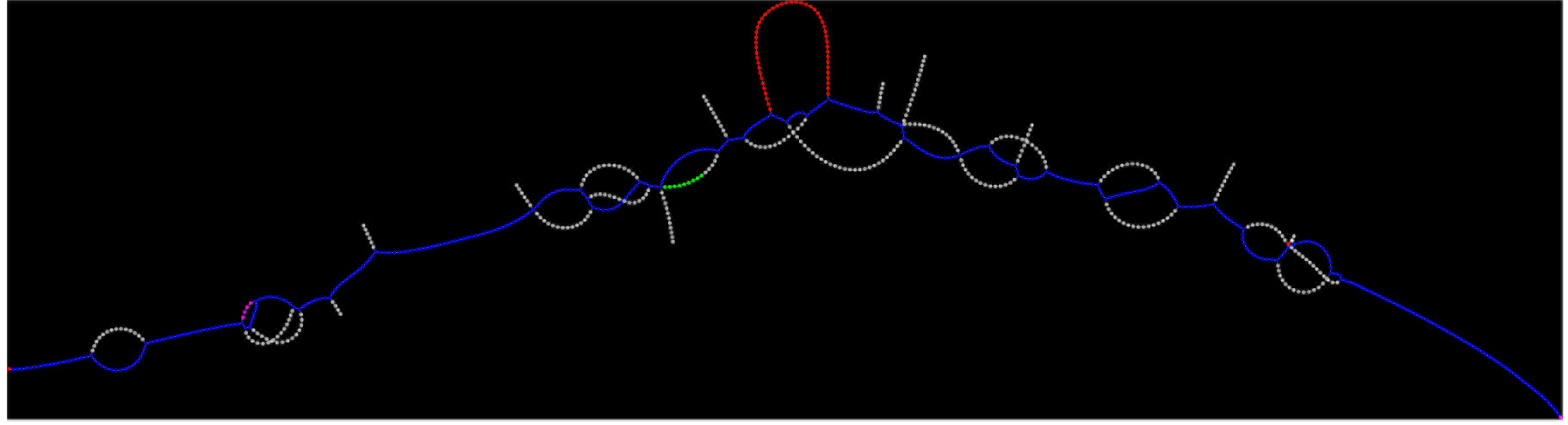
Colored DeBruijn illustration. Example of colored DeBruijn graph rendered using Lancet for a short region of 400bp containing an insertion. Blue nodes correspond to *k*-mers shared by both the tumor and the normal samples, red nodes correspond to *k*-mers private to the tumor, green nodes correspond to *k*-mers private to the normal, and white nodes correspond to low coverage *k*-mers due to sequencing errors.

We here introduce a new somatic SNV and indel caller, *Lancet*, which uses localized colored DeBruijn graphs (**Fig. 1**) to detect somatic variants with high accuracy in paired tumor and normal samples. Lancet builds upon the effective assembly engine we introduced in the Scalpel^2^ variant caller, that localizes the assembly to small genomic regions. However, unlike Scalpel, Lancet jointly assembles reads from a tumor and a matched normal sample into colored DeBruijn graphs that are automatically optimized according to the repeat composition of each sequence (**Supplementary Fig. 1** and **Online Methods**). The colored DeBruijn graph assembly paradigm was initially introduced and applied to detection and genotyping of both simple and complex germline variants in a single individual or population^3^. We here demonstrate that this paradigm is even more powerful in the context of somatic variant detection. Unlike the initial work of Iqbal *et al.*, where the colored DeBruijn graph is constructed for the whole genome, Lancet builds a local colored DeBruijn graph in a short genomic region (default 600bp) following the micro-assembly paradigm^1, 2^. The local assembly paradigm makes a very detailed analysis of the graph structure computationally tractable, allowing the detection of low frequency mutations private to the tumor without sacrificing the sensitivity to call longer mutations. In the Lancet framework, somatic variants correspond to simple paths in the graph whose nodes (*k*-mers) belong only to the tumor. Partially supported variants in the normal sample can be easily detected and classified as germline variants (**Supplementary Fig. 2**). Among its many features, Lancet employs: 1) an Edmonds-Karp style network-flow algorithm to efficiently enumerate all haplotypes in a genomic region; 2) on-the-fly short tandem repeat (STR) analysis of the sequence context around each variant; 3) a highly reliable scoring system; 4) carefully tuned filters to prioritize higher confidence somatic variants; and 5) a simple and efficient active region module to skip the analysis of genomic regions with no evidence of variation (**Online Methods**). Finally, in additional to running the tool in discovery mode, Lancet can be used interactively for an in-depth analysis of a region of interest, similarly to other bioinformatics utilities used for operating on BAM files, such as samtools^4^, bamtools^5^, bedtools^6^, etc. Colored DeBruijn graphs can be easily exported and rendered to visualize variants of interest in graph space (**Fig. 1**), which can help in confirming a variant. This feature complements read alignment visualization tools such as the Integrative Genomics Viewer (IGV)^7^ and provides another useful view into the data that supports variant calling.

## Results

We performed extensive experimental comparisons using several synthetic and real-world datasets designed to assess the variant calling abilities of Lancet under diverse tumor clonality/cellularity and sequencing conditions on a range of Illumina platforms (HiSeq 2000, HiSeq 2500, HiSeq X) commonly used for whole-genome sequencing. We compared Lancet to some of the most widely used somatic variant callers, including MuTect^8^, MuTect2, LoFreq^9^, Strelka^10^, and Strelka2^11^. Benchmarking datasets include (1) virtual tumors generated from real germline sequencing reads, that contain a predefined list of somatic mutations with known variant allele fractions (VAF); (2) synthetic tumors from the ICGC-TCGA DREAM mutation calling challenge^12^; (3) matched tumor and normal from a medulloblastoma case from the ICGC PedBrain Tumor project^13^; and (4) real data from a highly genetically concordant pair of primary and metastatic cancer lesions^14^.

**Figure 2.**
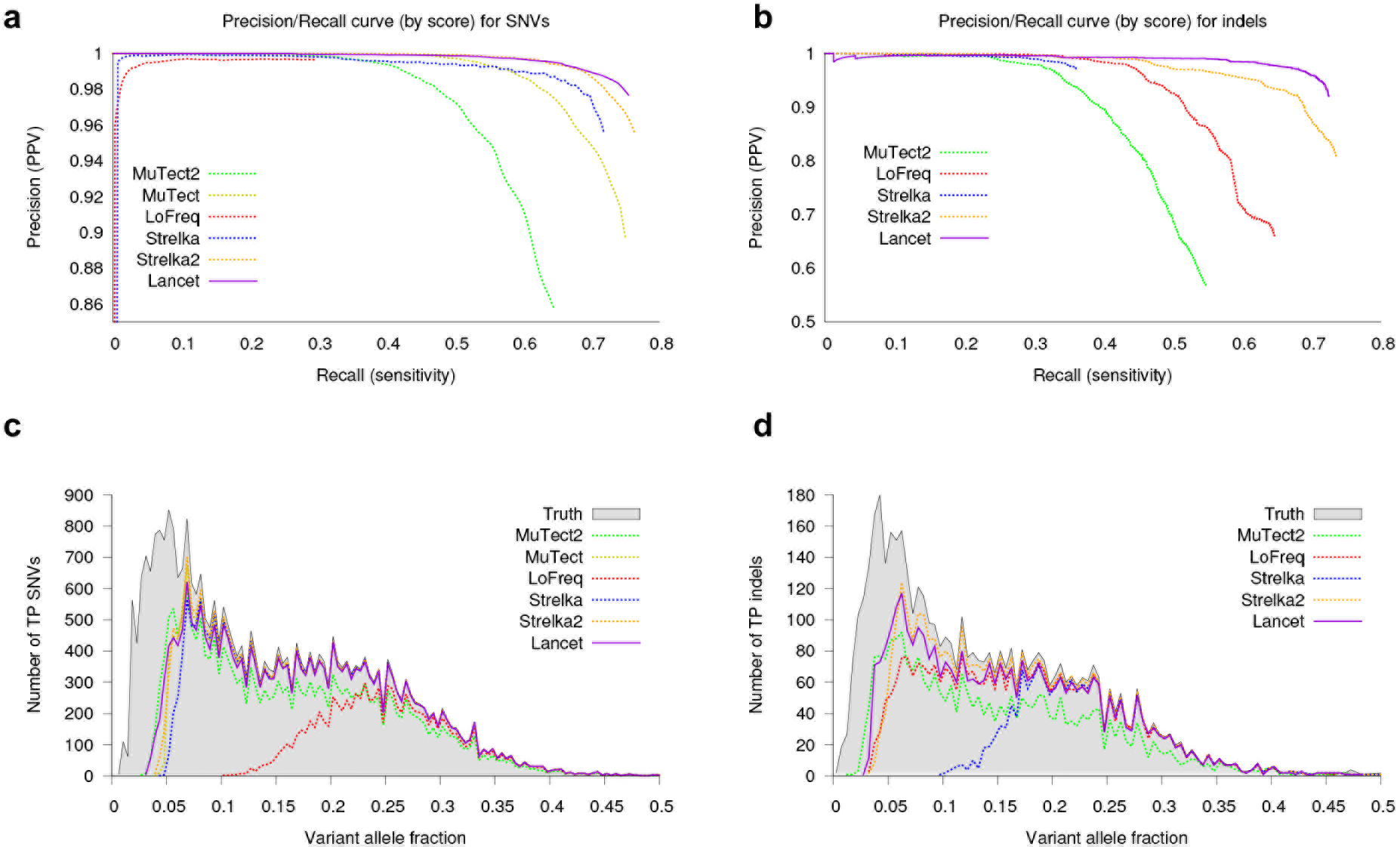
Performance of Lancet and other methods on the virtual tumors. (**a**) Precision/recall curves for somatic SNVs called by Lancet, MuTect, MuTect2, LoFreq, Strelka, and Strelka2 on the virtual tumor. Curves are generated by sorting the variants based on the confidence or quality score (QUAL) assigned by each tool. Each point on the curve corresponds to precision and recall of all the SNVs with confidence score less or equal to a specific quality threshold. The curve for an ideal tool (with no errors) should start from the top left corner (with precision=1) and produce a straight horizontal line. Any deviation from a straight line is due to errors introduced by the variant calling process. Specifically, deviations at low recall rates are indicative of low performance of the scoring system adopted by the tool (false positive variants reported with high score). (**b**) Precision/recall curves for somatic indels called by Lancet, MuTect2, LoFreq, Strelka, and Strelka2 on the virtual tumor. Number of true-positive (**c**) SNVs and (**d**) indels at different variant allele fractions for each method and for the truth call set.

### Virtual tumors

Using a strategy similar to the one described in the MuTect paper^8^, we generated virtual tumors by introducing reads that support real germline SNVs and indels in HapMap sample NA12892, from an unrelated HapMap sample NA12891, both sequenced on the Illumina HiSeq X system. Only actual sequencing data was used to spike-in somatic variants at a ladder of variant allele fractions at variable loci identified in those sample as part of the 1000 Genomes Project (**Supplementary Fig. 3** and **Online Methods**). By knowing the true somatic variants and controlling the VAF of inserted mutations, we use the virtual tumors to test the methods’ ability to call somatic mutations at predefined, including very low, VAFs. Precision/recall curves of somatic variant calls, sorted by their confidence score, show that Lancet outperforms all other somatic callers analyzed in this study on this dataset, especially for indels (**Fig. 2a-b**). On this dataset, Lancet behaves close to an (ideal) variant caller that makes no errors (straight line with precision=1) demonstrating a highly reliable scoring system for both SNVs and indels. The other tools tend to either introduce errors early by assigning high scores to false positive variants or substantially worsen in precision at higher recall rates. Although the truth set contains a handful of somatic STR mutations (**Supplementary Fig. 4**), analysis of indels called by each tool shows higher false positive rate of somatic STR indels for Strelka2, LoFreq, and MuTect2 compared to Lancet and Strelka (**Supplementary Fig. 5**); interestingly, the false positive STR indels are highly discordant across callers (**Supplementary Fig. 6b)**. When calling indels, Lancet and Strelka2 demonstrate higher sensitivity (**Supplementary Fig. 6a**) in particular for variants with VAF < 10% (**Fig. 2d**), however Lancet loses the least amount of precision compared to the other tools (**Fig. 2b**). All the callers show similar performance in the detection of indels with VAF>10%, with the exception of Strelka, whose sensitivity for indels is comparable to the other methods only at 20% VAF or above. Excluding LoFreq, all the tools show similar sensitivity to detect SNVs across the VAF spectrum (**Fig. 2c**), however Lancet’s superior accuracy is highlighted in the precision/recall curve (**Fig. 2a**). Finally, Lancet produces by far the best overall F_1_-score across all the tested methods on the virtual tumor for indel calling (**Tables 1** and **2**). Lancet and Strelka2 achieve the same F_1_-score on SNVs calling, however Lancet generates half the number of false positives compared to Strelka2. Analysis of the reference and alternative allele counts shows great variability in the number of supporting reads for each tool, due to the different methods and filters used in selecting the reads. As expected, most false positive indels have few reads containing the alternative allele; this is largely the case for Lancet, while other tools (e.g., MuTect2) also report false positives indels with higher support for the alternative allele, indicating a problem in selecting/filtering the set of alignments that support the mutations either in the tumor or the normal (**Supplementary Fig. 7**). Strelka has the lowest number of false positive calls but the distribution of supporting reads highlights its limited power in detecting indels with very low support.

**Table 1.** Somatic indel detection performance on the virtual tumor. Tools sorted in descending order of F_1_-score.

|  | # of calls | TP | FP | FN | Recall | Precision | FDR | F<sub>1</sub>score* | Max F<sub>1</sub>score† |
| --- | --- | --- | --- | --- | --- | --- | --- | --- | --- |
| **Lancet** | 3891 | 3586 | 305 | 1359 | 0.72 | 0.92 | 0.078 | **0.81** | **0.81** |
| **Strelka2** | 4514 | 3647 | 867 | 1298 | **0.73** | 0.81 | 0.192 | 0.77 | 0.78 |
| **LoFreq** | 4853 | 3210 | 1652 | 1744 | 0.64 | 0.66 | 0.340 | 0.65 | 0.67 |
| **MuTect2** | 4873 | 2712 | 2071 | 2233 | 0.54 | 0.56 | 0.432 | 0.55 | 0.58 |
| **Strelka** | 1846 | 1793 | 53 | 3152 | 0.36 | **0.97** | **0.028** | 0.52 | 0.71 |

\* F<sub>1</sub>score: harmonic mean of precision and recall, $2 \times (\text{precision} \times \text{recall}) / (\text{precision} + \text{recall})$ ; † Maximum F<sub>1</sub>score computed for each combination of precision and recall along the precision/recall curve.

**Table 2.** Somatic SNV detection performance on the virtual tumor. Tools sorted in descending order of F_1_-score.

|  | # of calls | TP | FP | FN | Recall | Precision | FDR | F<sub>1</sub>score* | Max F<sub>1</sub>score |
| --- | --- | --- | --- | --- | --- | --- | --- | --- | --- |
| **Lancet** | 24413 | 23848 | 565 | 7744 | 0.75 | 0.98 | 0.023 | **0.85** | **0.85** |
| **Strelka2** | 25249 | 24132 | 1117 | 7460 | **0.76** | 0.96 | 0.044 | **0.85** | **0.85** |
| **Strelka** | 23891 | 22741 | 1150 | 8851 | 0.72 | 0.95 | 0.048 | 0.82 | 0.82 |
| **MuTect** | 50228 | 23713 | 2792 | 7879 | 0.75 | 0.89 | 0.055 | 0.82 | 0.82 |
| **MuTect2** | 23779 | 20393 | 3386 | 11199 | 0.65 | 0.86 | 0.142 | 0.74 | 0.74 |
| **LoFreq** | 9404 | 9370 | 34 | 22222 | 0.3 | **0.99** | **0.003** | 0.46 | 0.46 |

\* F<sub>1</sub>score: harmonic mean of precision and recall, $2 \times (\text{precision} \times \text{recall}) / (\text{precision} + \text{recall})$ ; † Maximum F<sub>1</sub>score computed for each combination of precision and recall along the precision/recall curve.

### Synthetic tumors

We performed an additional comparison using the synthetic tumors from the ICGC-TCGA DREAM mutation calling challenge #4. This dataset was the most difficult to analyze due to a combination of complex clonality and cellularity of the tumor sample, which contained two sub-clones of 30% and 15% allelic fraction. Similarly to the virtual tumors, raw data from a deeply sequenced sample was randomly sampled into two non-overlapping subsets of equal size. Then a spectrum of mutations, some randomly selected and some targeting known cancer-associated genes, was introduced in one of the two samples (the tumor), using BAMSurgeon (https://github.com/adamewing/bamsurgeon). While somatic SNVs are spiked in by altering the original reads, in the case of indels synthetic reads containing the desired mutation were simulated and used to replace a fraction of the original reads from the same region. We discovered that the truth set for this dataset contains many variants with supporting reads coming only from one strand (thus introducing a strong strand bias), and for this experiment we turned off Lancet’s strand bias filter. In real tumors, such strong strand bias is unlikely to happen. Precision/recall curve analysis (**Fig. 3a**) together with the precision, FDR, and F_1_-score values (**Supplementary Tables 1** and **2**) show that on this dataset Lancet outperforms all other somatic callers for indel calling. As reported in previous studies^2, 15^, assembly based methods, such as Lancet and MuTect2, demonstrate substantially more power to detect indels of 50 base pairs or longer compared to alignment-based methods (**Fig. 3b**). Given the longer size range of indels spiked in this dataset, we also ran Streka2 in combination with Manta^16^, which is the recommended protocol for best somatic indel performance. This combination is indeed more sensitive to longer indels, but it is still subject to higher error rate compared to Lancet. Analysis of the size distribution of called variants outside of STRs shows that both MuTect2 and LoFreq have strong bias towards calling longer false positive indels (**Fig. 3c**). IGV inspection of a random subset of LoFreq calls on the ICGC-TCGA DREAM data highlights that the false positive indels are typically due to mis-alignment of the supporting reads in the normal (**Supplementary Fig. 8**). Most of the MuTect2 false positive insertions instead correspond to breakpoints of larger structural variants that are misinterpreted as small insertions (**Supplementary Fig. 9-10**). For SNV detection, Lancet shows comparable results to MuTect2, the best performing method for this dataset (**Supplementary Fig. 11**). Strelka2 shows an impressive precision/recall curve for SNVs up to 0.6 recall, however its precision drops considerably afterwards.

**Figure 3.**
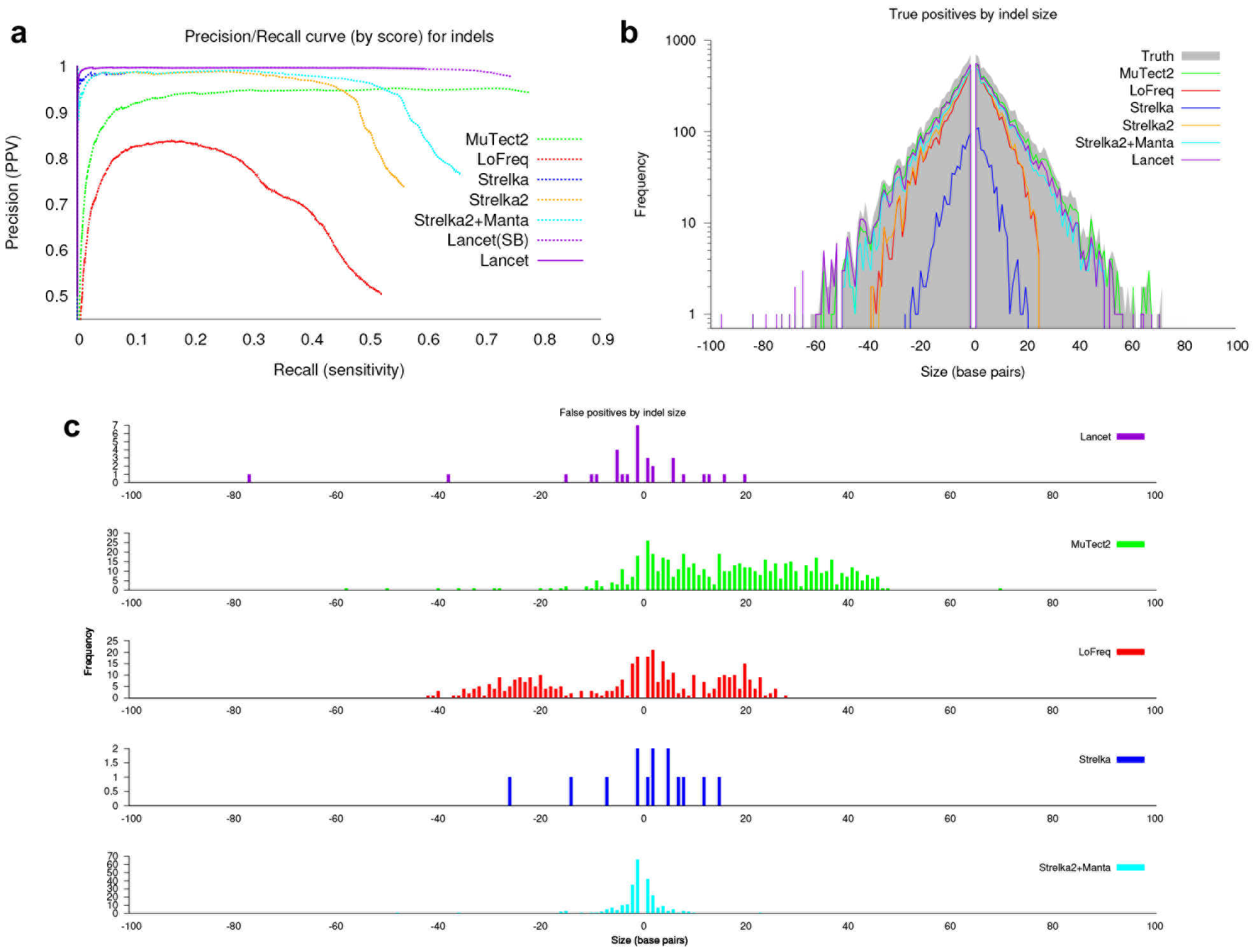
Indel performance of Lancet and other methods on the synthetic tumor #4 of the ICGC-TCGA DREAM mutation calling challenge. (**a**) Precision/recall curve analysis of somatic indels called by Lancet, MuTect, MuTect2, LoFreq, Strelka, Strelka2, and Strelka2+Manta. Lancet^SB^ is the version of Lancet run with strand bias filter turned off. (**b**) Size distribution of true positive indels for each method. Assembly based methods (Lancet, MuTect2, and Strelka2+Manta) demonstrate substantially more power to detect longer indels, while alignment-based methods (LoFreq, Strelka, and Strelka2) have reduced power to detect larger mutations, in particular insertions. (**c**) Size distribution of false-positive indels, excluding STRs, plotted separately for each method. LoFreq false positive indels are mostly due to mis-alignment of the reads supporting the indel in the normal, while most of the MuTect2 false positive insertions instead correspond to breakpoints of larger structural variants (e.g., inversion, translocations) that are misinterpreted as insertions. Lancet, Strelka and Strelka2 show the lowest number of false positives although Lancet has superior sensitivity compared to Strelka and Strelka2+Manta on this dataset.

### Normal tissue/tumor pair

We next analyzed real data from a case of medulloblastoma used in the cross-centers benchmarking exercise of the International Cancer Genome Consortium (ICGC)^13^. Unlike the synthetic tumors of the ICGC-TCGA DREAM mutation calling challenge, no single mutation was spiked-in, but rather a curated list of somatic mutations (SNVs and indels) was compiled (the Gold Set). Due to the heterogeneity of the raw data (multiple library protocols, Illumina sequencers, read lengths, and fragment sizes), this dataset is particularly noisy and challenging to analyze. Moreover, differently from the previous datasets used in this study, the majority of indel calls contained in the Gold Set are located within STRs (**Supplementary Fig. 13a**). Variant calling accuracy of all tools is generally inferior in comparison to the previous benchmarking experiments (**Supplementary Fig. 12**) but final precision and recall values are in agreement with the results reported by the ICGC benchmarking team. Strelka2 and LoFreq have better precison/recall curves for indels up to 0.5 recall, but Lancet shows the best final trade-off between precision and recall (F_1_-score) and it ranks second in SNV detection, after LoFreq (**Supplementary Tables 3** and **4**). Although LoFreq and Strelka2 have higher indel recall rates (**Supplementary Tables 3** and **Supplementary Fig. 13b**), their final precision is substantially lower compared to Lancet and Strelka (**Supplementary Fig. 13c**), indicating that these tools may have difficulties in handling the noise in the data. Inspection of the F_1_-score values, as a function of recall, shows all callers favor sensitivity over specificity in this dataset (**Supplementary Fig. 14**) - indicating that they have likely been optimized for higher quality data. As is the case with virtual tumors, false positive indels within STRs are highly discordant across callers in the medulloblastoma dataset (**Supplementary Fig. 13c-d**), thus confirming an overall lower quality of these calls. In contrast, Lancet reports a very small number of false positive indels without losing sensitivity (**Supplementary Fig. 13b-c**).

### Normal tissue/primary tumor/metastasis trio

Finally, we analyzed a pair of highly genetically concordant primary and metastatic cancer lesions to check the robustness of different methods to identify shared and private somatic mutations. Concordance of SNVs shared between the primary and metastasis is much higher compared to indels among the analyzed tools, however higher agreement of the called indels is achieved when indels within STRs are removed (**Supplementary Fig. 15**). These results once more highlight the problem of detecting somatic STRs and emphasize the challenging, but necessary, task of integrating indel calls across different methods.

## Discussion

Across the four datasets analyzed in this study, we discovered that the major source of disagreement between callers originates from somatic variants called within STRs, in particular if the motif is two base pairs or longer. Moreover, Venn diagram analysis shows substantial disagreement between the callers for the false positive somatic STR calls. Since the virtual tumors were created by partitioning the raw reads from a single real sample, we infer that the erroneous STR indels are the results of higher replication slippage at those sites that most tools misclassify as somatic events. In contrast, thanks to reliable scoring and filtering systems and the employment of the local assembly engine, Lancet makes fewer errors at STR sites. Alignment based tools, such as LoFreq, are inherently more prone to misclassify longer variants as somatic. Lancet instead natively corrects for mis-aligned reads thanks to the joint assembly of the tumor and normal reads in the same colored DeBruijn data structure, which also provides more precise estimation of the variant allele fraction. Our extensive comparative analysis also indicates that somatic callers are now optimized for higher quality data, although inspection of the max F_1_-score values suggests that better performance is achievable on noisy data with more stringent quality cutoffs.

The key novel feature introduced by Lancet is the usage of colored DeBruijn graphs to jointly analyze tumor and normal reads. This strategy substantially increases the accuracy of identifying mutations, especially indels, private to the tumor. Precision/recall curve analysis demonstrates that Lancet has a reliable variant quality scoring system, which is critical for prioritizing somatic variants. Lancet shows high precision when calling somatic mutations and provides robust calls across data generated by different Illumina sequencers. Due to its pure local-assembly strategy, Lancet currently has longer runtimes compared to alignment based methods (**Supplementary Table 5**), which is an area we plan to improve upon in the future releases of the tool. In addition to being used as a genome-wide analysis tool, Lancet can be used interactively to call variants and render colored DeBruijn graphs at small genomic regions of interest. In summary, Lancet provides highly accurate genome-wide somatic variant calling of SNVs and indels, and, given all its new features, we anticipate Lancet to become an invaluable resource for the bioinformatics community working on cancer.

## Methods

### Lancet workflow

Lancet uses the same local assembly engine initially developed for the Scalpel variant caller^2^ but it introduces many new features specifically designed for somatic analysis of tumor and matched normal next-generation sequencing data. The algorithm starts by decomposing the whole genome into overlapping windows of a few hundred base pairs (600bp by default). Each region is then locally assembled, except repetitive regions that have an excessive number of mapped reads (default 10,000), using the workflow depicted in **Supplementary Fig. 1**. Reads mapping within each region are extracted from the tumor and normal BAM files and decomposed into *k*-mers which are then used to build a colored DeBruijn graph as described in section “Colored DeBruijn graph construction”. Reads used for the assembly are carefully selected to reduce the number of possible artifacts in the graph that could confound variant detection. The details of the read selection process and the various filters applied are described in section “Read selection”. The graph is initially built using a small k-mer value (starting with a default of *k* = 11) which allows incorporation of reads supporting very low coverage variants. However, the k-mer parameter is automatically increased along the scale of odd numbers, to avoid the presence of perfect and near perfect repeats (default up to 2 mismatches) in the graph that can confound variant detection by introducing false bubbles, described in section “Repeat analysis”. The graph complexity is then reduced by removing low-coverage nodes, dead-ends, short-links, and by compressing chains of uniquely linked nodes (section “Graph cleanup”). Once a repeat-free graph has been constructed, it is anchored to the reference by selecting one *source* and one *sink* node corresponding to unique *k*-mers located within the current window. All possible *source*-to-*sink* paths are then efficiently enumerated using an Edmonds–Karp style algorithm described in section “Paths enumeration”. The assembled sequences from each path are aligned to the reference window using a sensitive Smith-Waterman-Gotoh alignment algorithm with affine-gap penalties. Finally, the alignments are parsed to extract the signature of different mutations (single nucleotide variant, insertion, and deletion).

### Read selection

Reads aligning to the genome are extracted from the tumor and normal BAM files and used for local assembly with the exception of the following set of reads. (1) PCR duplicates marked using the Picard MarkDuplicates module (https://broadinstitute.github.io/picard) – removing PCR duplicates is necessary to correctly estimate coverage and support for variant calls. (2) Reads aligned with low mapping quality (< MP, default 15) – reads with low mapping quality may be mapped to the wrong genomic location or aligned with incorrect signature. (3) Reads which are highly likely to be multi-mapped. Depending on which version of the BWA aligner is employed, there are two ways to identify these reads. In the case of BWA-MEM, multi-mapped reads are assigned equal values in the AS and XS tags, however we slightly relaxed this constraint to identify reads which are highly likely to be multi-mapped (|AS-XS| ≤ δ where δ =5). If BWA-ALN is employed, multi-mapped reads are marked using the XT:Z::R tag, nonetheless, their mapping quality is not necessarily zero. This is because mapping quality is computed for the read pair, while XT is only determined from a single read. For example, when the mate of a read can be mapped unambiguously, the read can still be mapped confidently and thus assigned a high mapping quality. In addition to the XT tag, multi-mapped reads are also identified using the XA tag which is used to list the alternative hits of the read across the genome. Finally, to maximize the sensitivity to detect variants that are also present in the normal sample, no filter is applied when extracting the reads aligned to the normal.

### Colored DeBruijn graph construction

The key data structure used by Lancet is the colored DeBruijn graph constructed using the reads from both the tumor and the matched normal samples. **Fig. 1** shows an example of the DeBruijn graphs generated by Lancet. Formally the graph is defined as *G* (*V, E, C*) where *V* is the set of vertices/nodes corresponding to the different *k*-mers extracted from the reads, *E* is the set of edges connecting two nodes having a k-1 perfect match between their respective *k*-mers, and *C* is the coloring scheme (labels) used to indicate whether the k-mer has been extracted from the tumor or normal sample. To account for the double-strandedness of DNA, Lancet constructs a bi-directed DeBruijn graph where each node stores both forward and reverse complement of each *k*-mer. The graph is augmented with ancillary information extracted from the raw sequencing data, specifically each node stores (*i*) the *k*-mer counts split by strand, (*ii*) the list of reads where the *k*-mers were found, and (*iii*) the Phred quality for each base. The *k*-mers from the reference sequence are also extracted and incorporated into the graph. Sequencing data is typically generated from short-insert paired-end DNA libraries and the variable fragment size distribution can sometimes cause two paired reads to overlap each other. Therefore, coverage needs to be adjusted to avoid over counting the overlapping portion of the two reads. This is easily accomplished in the DeBruijn graph framework since *k*-mers extracted from the overlapping segment come from reads that share the same query template (QNAME) in the BAM file. If this condition is detected, the k-mer count is adjusted to only count one copy of the two *k*-mers.

### Graph cleanup

Sequencing errors, coverage fluctuations, and mapping errors increase the graph complexity by introducing nodes and edges that confound the analysis. Lancet utilizes several graph operations and transformations designed to remove spurious nodes and edges introduced during graph construction. First, low-coverage nodes, which are typically associated with sequencing errors, are removed if the corresponding *k*-mer count is below a specific user defined threshold (default 1) or if the coverage ratio is below a certain user defined value (default 0.01). Second, dead-ends are removed, which present themselves as a sequence of uniquely linked nodes that do not connect back to the graph (also called short tips). Dead-ends formed by *n* (default 11) or more nodes are removed from the graph. Next *short-links* are removed, which are short connections composed by fewer nodes than theoretically possible given the k-mer value used to build the graph. **Supplementary Figure 16** illustrates one exemplary short-link scenario. This type of connection is typically due to sequence homology between closely located repeats (e.g., Alu repeats), but it can also happen in the case of long homopolymers, and other short tandem repeats, where the tandem repetition of the motif can result in the construction of a tiny bubble in the presence of a heterozygous mutation. Those tiny bubbles need to be kept in the graph as they may represent true variation, while short-links like the one depicted in **Supplementary Fig. 18** can be safely removed. Therefore, connections at non-STR sites formed by *m* (<< *k*) or less nodes and whose minimum coverage node is *c* < √*c*_*avg*_ are removed from the graph, where *c*_*avg*_ is the average coverage across the window. Finally, the graph is compressed by merging chains of uniquely linked nodes into super nodes.

### Repeat analysis

Small scale repeats are a major challenge for accurate variant calling, specifically for indels^1^. To avoid introducing errors at those loci, Lancet employs the same repeat analysis procedure introduced in the Scalpel algorithm. Specifically, the sequence composition in each window is analyzed for the presence of perfect or near-perfect repeats (up to a specified number of mismatches, 2 by default) of size *k*. Similarly, the graph is inspected for the presence of cycles (perfect repeats) or near-perfect repeats in any of the source-to-sink paths. If a repeat structure is detected, a larger *k*-mer value is selected and the repeat analysis is performed again on both the reference sequence and the newly constructed graph, until a repeat-free graph is constructed or the *k*-mer size has reached a maximum value (101 by default). To avoid using *k*-mers which are reverse complement of their own sequences, only odd values of *k* are used to build the graph. This iterative strategy is a key feature of the Lancet algorithm which automatically selects the optimal *k-* mer size according to the sequence composition of each genomic window.

### Paths enumeration

Enumerating all possible haplotypes can take time, growing exponentially with the number of bubbles present in the graph. To reduce the computational requirements of the graph traversal down to polynomial time, we employ an Edmonds–Karp style algorithm for fast enumeration of all possible haplotypes. The idea behind the algorithm is to find the minimum number of paths from source to sink that cover every edge in the graph (edge and nodes can be visited more than once). The pseudo code of the algorithm is presented below. Since every node is visited (possibly multiple times), it is easy to show that, although the same variant could be discovered multiple times, no variant is missed from the analysis. Straightforward complexity analysis of the pseudocode shows that the worst-case time complexity is O(*E*^*2*^+*EV*): at least one edge is visited at each iteration (step 5) accounting for O(*E*) time, and each call to the graph traversal (step 2) takes O(*E*+*V*) where *E* is the number of edges and *V* the number of nodes in the graph. As such, a trivial upper bound for the whole procedure is O(*E*) × O(*E*+*V*) = O(*E*^*2*^+*EV*).

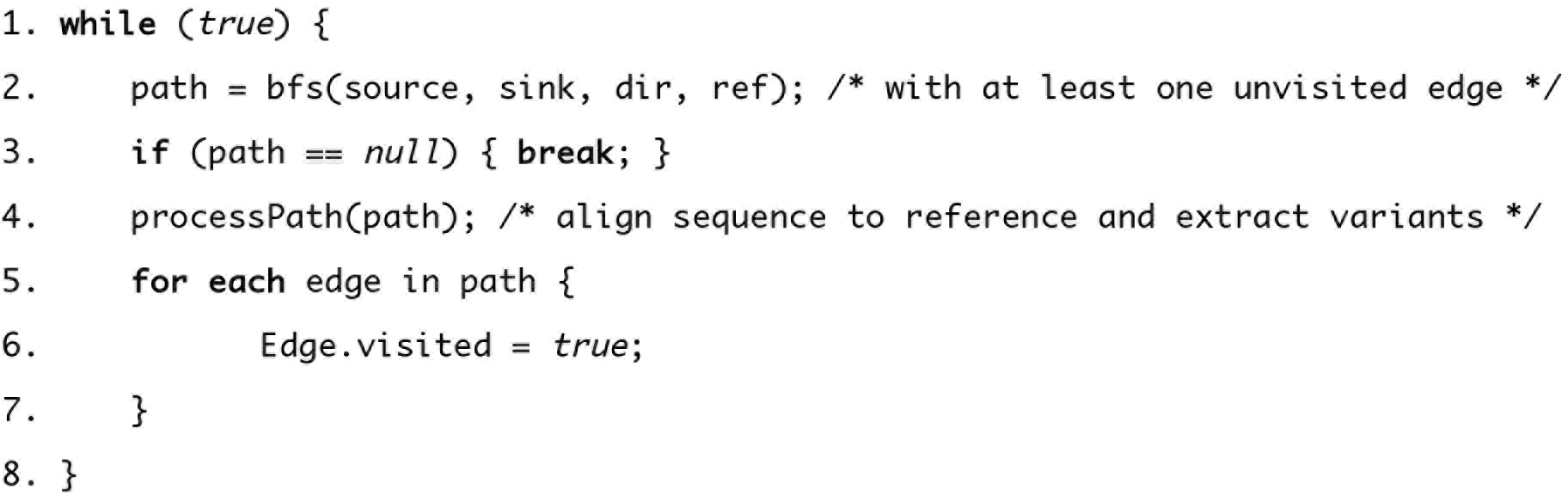

### Active regions

The idea behind the active region module is to avoid wasting time processing (read extraction, local assembly, re-alignment) regions without evidence for variation. Regions where all reads map to the reference without any mismatches can be trivially discarded. However, the error rate of the Illumina sequencing technology (~0.1 percent), in combination with high coverage, makes the scenario of alignments with no mismatches in a region very unlikely. The policy adopted by Lancet is to consider a region as “active”, either in the tumor or the normal sample, if a minimum of *N* (aligned) reads support a mismatch, indel, or soft-clipped sequence at the same locus (**Supplementary Fig. 17**), where *N* is equal to the minimum alternative count support specified for somatic variants (3 by default). This policy is implemented on the fly by simple and fast parsing of the MD and CIGAR strings. This step is functionally similar to the active region module employed in MuTect and other tools, however Lancet follows a pure assembly approach, where all variant types (SNVs, insertions and deletions) are detected through local assembly. When tested on an 80x/40x coverage pair of tumor/normal samples sequenced with 150bp reads, Lancet’s active region strategy discards on average between ~10% and ~20% of the total number of windows. However, due to its pure assembly strategy, Lancet typically requires higher runtimes compared to the hybrid approach employed by MuTect2 and Strelka2 (**Supplementary Table 5**). To achieve faster runtimes and to discard more windows, the parameter *N* can be increased when analyzing samples sequenced at coverage higher than 80x/40x.

### Scoring variants

Fisher’s Exact test is used to determine if a mutation has non-random associations between the allele counts in the tumor and in the normal samples. Specifically, given a somatic mutation, reference and alternative reads supporting the variant both in the tumor and the normal are collected and stored into a 2-by-2 contingency table which is then used to compute a Phred-scaled Fisher’s exact test score, *S*_*(fet)*_, according to the following formula:

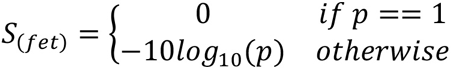

where *p* is the exact probability of the 2-by-2 contingency table given by the hypergeometric distribution.

### Variant filters

Lancet generates the list of mutations in VCF format^17^ (v4.1). All variants (SNVs and indels) either shared, specific to the tumor, or specific to the normal are exported as part of the output. Following the VCF format best practices, high quality variants are labelled as PASS in the FILTER column. Several standard filters, all of which have tunable parameters, are applied to remove germline calls and low quality somatic variants as describe here:

1. *Low/high coverage:* mutations located in substantially low coverage regions of the normal (default < 10) or tumor (default < 4) are removed since there is a high chance for coverage bias towards one of the alleles.
2. *Variant allele fraction*: mutations characterized by a very low variant allele fraction in the tumor (default < 0.04) are filtered because they are likely to be false positive calls. Likewise, variants whose variant allele fraction is high in the normal (default > 0.0) are considered to be germline calls.
3. *Alternative allele count*: analogously to the allele fraction filter, mutations with low alternative allele count (default < 3) in the tumor are likely to be false positive calls and are flagged as low quality. While variants with a high alternative allele count in the normal (default > 0) are considerate to be germline mutations.
4. *Fisher's exact test (FET) score*: mutations with a very low FET score are flagged as low quality. Due to their inherently different error profiles, separate thresholds are used for non-STR variants (default < 5.0) and STR variants (default < 25.0).
5. *Strand bias:* this filter rejects variants where the number of alternative counts in the forward or reverse strand is below a certain threshold (default < 1).
6. *Microsatellite*: microsatellites (or short tandem repeats) are highly mutable genetic elements subject to high rate of replication slippage events (especially homopolymers), which reduces variant callers’ ability to distinguish between sequencing errors and true mutations. As such, mutations located within microsatellites or in their proximity (default 1 base pair away) are recognized and flagged by Lancet. By default, microsatellites are defined as sequences composed of at least 7bp (total length), where the repeat sequence is between 1bp and 4bp, and is repeated at least 3 times. The user can adjust these parameters to define any type of microsatellite motif size and length as required by different applications.

### Read alignment and BAM file generation

Sequencing reads were aligned to the human reference hg19 using BWA-MEM (v.0.7.8-r455) with default parameters. Alignments were converted from SAM format to sorted and indexed BAM files with SAMtools (v.1.1). GATK software tools (v.2.7-4) were used for improving alignments around indels (GATK IndelRealigner) and base quality recalibration (GATK base quality recalibration tool) using recommended parameters. Finally, the Picard tool set (v.1.119) was used to remove duplicate reads. The final BAM files generated by this process were used as input for all the variant callers used in this study.

### Virtual tumors

We created virtual tumors using a strategy similar to what was employed in the MuTect paper^8^. We sequenced HapMap sample NA12892 at high coverage on the Illumina HiSeq X system using PCR-free protocol and partitioned the set of reads into two groups of 80x and 40x average coverage to use as tumor and normal respectively. Reads were mapped using the alignment procedure described in section “Read alignment and BAM file generation”. We then used an unrelated HapMap sample NA12891 sequenced on the same Illumina HiSeq X system to introduce realistic SNVs and indels by swapping a predefined number of reads between the two samples at loci where NA12892 is homozygous reference and NA12891 is homozygous variant (**Supplementary Fig. 3**). The list of selected loci is based on the 1000 Genomes Project phase 3^18^ call set and the number *N* of reads that were swapped between the two samples followed a binomial distribution with mean μ = 0.05, 0.1, 0.2, 0.3. This procedure allowed us to spike-in realistic mutations with known variant allele fractions, but the length of indels was limited by the short size range currently included in the 1000 Genomes call set. Specifically, the longest insertion and deletions that we were able to spike in were 13 bp and 35 bp respectively. We used this process separately for SNVs and indels to create two pairs of tumor/normal samples with 31,592 somatic SNVs and 4,945 somatic indels respectively. The virtual tumor BAM files together with the list of true variants are freely available for download at the New York Genome Center ftp site (ftp://ftp.nygenome.org/lancet).

### ICGC medulloblastoma benchmarking data

We downloaded the full set of FastQ files of the medulloblastoma patient (accession number EGAD00001001859) from the European Genome-phenome Archive (EGA, https://www.ebi.ac.uk/ega). The raw reads were generated by five different sequencing centers reaching a cumulative coverage of ~300X for both the tumor and the normal samples. We merged the raw FastQ files separately for the tumor and the normal samples and then aligned the reads using the alignment pipeline describe in section “Read alignment and BAM file generation”. Then we down-sampled the ~300X BAM files down to ~80X and ~40x for the tumor and the normal respectively using the Picard DownsampleSam module. The down sampled BAM files generated by this process were then used as input for all the somatic variant callers used in this study.

### Primary and metastatic cancer lesions data

Sequencing data for the paired primary and metastatic cancer lesions are publically available through the database of Genotypes and Phenotypes (dbGaP, https://www.ncbi.nlm.nih.gov/gap) with accession number phs000790.v1.p1. The same data is also available through the Memorial Sloan Kettering Cancer Center cBioPortal for Cancer Genomics (study “Colorectal Adenocarcinoma Triplets”). In this study, we used the sequencing data for sample EV-014 and the BAM files were created following the same procedure described in section “Read alignment and BAM file generation”, with the only difference that the normal, primary and metastatic samples have been realigned together (with GATK IndelRealigner) to further improve alignments around indels.

### Variant calling

We tested the variant calling abilities of eight different somatic variant callers: Lancet (v1.0.0), MuTect (v1.1.7), MuTect2 (v2.3.5), LoFreq (v2.1.2), Strelka (v1.0.14), Strelka2 (v2.8.3), Scalpel^2^ (v0.5.3), and VarDict^19^ (v328e00a). Although a larger number of somatic variant callers is available in the literature, we chose to compare Lancet against these methods because they are some of the most widely used approaches specifically designed for whole genome tumor/normal variant calling and they represent a combination of both assembly and alignment based methods. Default parameters were used for each tool. Results on the virtual tumors revealed Scalpel and VarDict to be outliers in terms of specificity (**Supplementary Fig. 18**), so we decided to exclude these two tools from the overall benchmarking experiments.

### Benchmarking workflow

We used the following procedure to perform the Precision/Recall curve analysis employed in this study:

1. First, we ran each tool with default parameters, as reported in the “Variant calling” section.
2. We kept only the PASS somatic variants within the autosomes together with chromosomes X, Y and sorted the variant calls, from highest quality to the lowest, according to the quality score reported by each method in the final VCF file (“FisherScore” for Lancet, “SomaticEVS” for Strelka2, “QSI” for Strelka, “QUAL” for LoFreq, “TLOD” for MuTect and MuTect2).
3. Due to the possibly ambiguous representation of indels around microsatellites and other simple repeats, we left normalized all the indels.
4. When comparing calls to the truth set or across the different methods, we matched two variants (SNV or indels) if they shared the same genomic coordinates (chromosome and start position) as well as if they have the exact same sequences (both in size and base pair composition) in the reference and alternative alleles.
5. Precision/recall values along the curve are then computed for each tool by processing the somatic calls in the sorted order generated in step 2.

### Code availability and system requirements

Lancet is written in C/C++ and is freely available for academic and non-commercial research purposes as an open-source software project at https://github.com/nygenome/lancet. Lancet employs two widely used next-generations sequencing analysis APIs/libraries, BamTools (https://github.com/pezmaster31/bamtools) and HTSlib (http://www.htslib.org/), to read and parse the information in the BAM file, which are included in the code distribution. The source code has no dependencies and it is easy to compile and runs across different operating systems (Linux and Mac OSX). Lancet supports native multithreading via pthreads parallelization. Analysis of one whole-genome (80x/40x) tumor-normal pair sequenced with 150 base pair reads usually requires 3000 core hours and a minimum of 20 GB of RAM on a modern machine after splitting the analysis by chromosome.

### Data availability

Data used in this study was retrieved from the 1,000 Genomes website (http://www.1000genomes.org), the European Genome-phenome Archive (EGA, https://www.ebi.ac.uk/ega<u>) with accession number</u> EGAD00001001859, the database of Genotypes and Phenotypes (dbGaP, https://www.ncbi.nlm.nih.gov/gap) with accession number phs000790.v1.p1, and the International Cancer Genome Consortium (ICGC, http://icgc.org/). The virtual tumors generated and analyzed in this study are freely available for download at the New York Genome Center public ftp site (ftp://ftp.nygenome.org/lancet).

## Acknowledgments

We thank Michael C. Schatz and Wayne Clarke for the helpful discussion and comments on the manuscript.

## Authors contributions

G.N. designed the algorithms, developed the software, and conducted the computational experiments. A.C., A.K.E., V.V., M.Z. contributed to the design of algorithms. K.A., E.A.B., M.S., assisted with benchmarking the different somatic variant caller. R.M. assisted with the integration of multiple high-throughput sequencing APIs. All authors assisted with the design and interpretation of the comparative analysis between the different methods. G.N. wrote the manuscript with input from all the authors. All of the authors have read and approved the final manuscript.

